# Characterizing in vivo degradation of electrospun biodegradable nanofibers by size-exclusion chromatography

**DOI:** 10.64898/2026.05.10.724172

**Authors:** Shingo Kunioka, Takumi Yoshida, Daisuke Naruse, Yuki Setogawa, Hiroyuki Miyamoto, Ryohei Ushioda, Yuta Kikuchi, Masahiro Tsutsui, Hiroyuki Kamiya, Kyohei Oyama

**Affiliations:** Department of Cardiac Surgery, Asahikawa Medical University, Asahikawa, Japan; Life Materials Development Section, Human Life Technology Research Institute, Toyama Industrial Technology Research and Development Center; Business Development section, Business Development and Quality Control Department, IAAZAJ HOLDINGS CO., LTD; Department of Cardiothoracic Surgery, Thomas Jefferson University Hospital, Philadelphia, PA, USA

## Abstract

Biodegradable electrospun nanofiber (NF) scaffolds have emerged as promising materials for tissue engineering applications, including vascular grafts, because their mechanical properties and degradability can be tuned. However, their in vivo degradation behavior remains poorly understood. In this study, we characterized the in vivo degradation profiles of representative biodegradable NF materials widely used in small-caliber vascular graft research, namely polycaprolactone (PCL), poly(D,L-lactide) (PLA), polyglycolic acid (PGA), and a PCL/PLA blend, by monitoring molecular weight changes in subcutaneous and vascular environments.

Electrospun NF sheets were implanted subcutaneously in mice, and tubular NF grafts were implanted into the abdominal aorta of rats. Samples were harvested for up to 48 weeks after implantation and analyzed primarily by size-exclusion chromatography (SEC) to assess time-dependent changes in molecular weight. Scanning electron microscopy (SEM) and solid-state ^13^C nuclear magnetic resonance (NMR) were additionally performed to evaluate ultrastructural and chemical changes associated with degradation.

SEC analysis revealed distinct material-specific degradation patterns. PCL showed the slowest degradation and retained a relatively high weight-average molecular weight (Mw) in both environments. PLA exhibited marked environment dependence, with near-complete degradation in the subcutaneous environment by 48 weeks, whereas scaffold structure was maintained in the vascular environment. The PCL/PLA blend showed earlier reduction in the high-molecular-weight fraction than PCL, indicating faster scaffold breakdown. PGA degraded most rapidly and could not be evaluated beyond 2 weeks in the subcutaneous model or in the vascular model because of early graft rupture. SEM analysis further demonstrated that progressive loss of fibrous ultrastructure over time was a common feature across all materials. In addition, NF scaffolds became resistant to organic solvent after implantation in vivo, and solid-state ^13^C NMR analysis of the solvent-insoluble fractions detected polymer-derived signals together with additional signals consistent with biological constituents.

These findings indicate that in vivo degradation of biodegradable NF scaffolds is material dependent, environment dependent, and more complex than simple hydrolytic chain cleavage alone. This study provides a quantitative framework for evaluating NF degradability and offers new insight into the design of biodegradable vascular grafts.

**Highlights:**

- SEC quantified long-term in vivo degradation of PCL, PLA, PGA, and PCL/PLA.
- Degradation was both material dependent and implantation environment dependent.
- In vivo nanofiber degradation involved structural and chemical changes beyond hydrolysis.

## 1. Introduction

With vascular diseases increasing worldwide, coronary artery bypass grafting and peripheral arterial bypass surgery remain the gold-standard treatments for small-caliber revascularization[1-4]. However, the current dependence on autologous vessels remains a major limitation, as suitable grafts are not always available in patients with prior vein harvest, diffuse vascular disease, poor vessel quality, or multiple revascularization requirements[5-9]. This clinical need has driven the development of clinically applicable small-caliber synthetic vascular grafts[10-13].

Biodegradable electrospun nanofiber (NF) scaffolds are considered a promising option for small-caliber vascular grafts because they can mimic the extracellular matrix while allowing control over mechanical properties and degradation rates. If scaffold degradation can be appropriately synchronized with host-tissue remodeling, NF-based grafts may enable gradual replacement of the implanted material by autologous tissue and thereby reduce long-term problems such as compliance mismatch[14-17]. A variety of biodegradable polymers have been used for this purpose, including polycaprolactone (PCL), poly(D,L-lactide) (PLA), polyglycolic acid (PGA), and blended materials such as PCL/PLA[18]. However, although these materials are widely used, their in vivo degradation behavior remains incompletely characterized. Most previous studies describe only approximate degradation timelines, while the actual degradation process, including differences among materials and implantation environments, remains poorly understood. A clearer understanding of the degradation behavior of commonly used NF materials would therefore be valuable for the rational design of NF-based vascular grafts, particularly because quantitative evaluation of degradation kinetics may help identify materials suited to different remodeling time courses and structural demands.

Size-exclusion chromatography (SEC) is an analytical technique that separates polymer molecules according to their hydrodynamic size and enables quantitative evaluation of molecular weight distribution. Because biodegradable NF scaffolds are degraded primarily through hydrolytic cleavage of polymer chains, SEC provides a useful approach for characterizing their in vivo degradation behavior through changes in molecular weight. Using this method, we aimed to characterize the in vivo degradation profiles of representative biodegradable NF materials widely used in small-caliber vascular graft research, namely PCL, PLA, PCL/PLA, and PGA, and to quantitatively compare their degradation behavior in subcutaneous and vascular environments.

## 2. Materials and Methods

### 2.1. Preparation of Electrospun Nanofiber Scaffolds

Electrospun nanofibers composed of polycaprolactone (PCL), poly(D,L-lactide) (PLA), polyglycolic acid (PGA), and a PCL/PLA blend were prepared under the following conditions. PCL nanofibers were electrospun from a 10% (w/w) PCL solution (Mw = 80,000; Sigma-Aldrich) dissolved in N,N-dimethylformamide (DMF) and tetrahydrofuran (THF) (3:7, w/w). Electrospinning was performed at 20 kV with a 20 cm tip-to-collector distance. PLA nanofibers were prepared from PLA (Mw = 135,000; Musashino Chemical Laboratory; L:D ratio = 85:15) dissolved in DMF and methyl ethyl ketone (MEK) (6:4, w/w) at a resin concentration of 16.7% (w/w). Electrospinning was performed at 20 kV with a 20 cm tip-to-collector distance. PCL/PLA nanofibers were prepared from a blended solution of PCL and PLA at a weight ratio of 8:2, dissolved in N,N-dimethylformamide (DMF), tetrahydrofuran (THF), and methyl ethyl ketone (MEK) (3.5:6:0.5, w/w) at a resin concentration of 11% (w/w). Electrospinning was performed at 20 kV with a 20 cm tip-to-collector distance. PGA nanofibers were prepared by dissolving PGA (intrinsic viscosity: 1.0-2.0; Polysciences Inc.) in 1,1,1,3,3,3-hexafluoro-2-propanol (HFIP) at a resin concentration of 5% (w/w). Electrospinning was performed at 10 kV with a 10 cm tip-to-collector distance.

For subcutaneous implantation, the electrospun nanofiber (NF) sheets were trimmed into 40 × 40 × 0.6 mm pieces. For vascular graft preparation, the NF sheets with a thickness of 30 µm were cut into 20-mm-wide strips, rolled onto a 2-mm polytetrafluoroethylene (PTFE) rod to achieve a wall thickness of approximately 500 µm, and sectioned into lengths of 5-7 mm, as described in our previous studies[9,19].

### 2.2. In Vivo Implantation Models (Subcutaneous and Vascular)

All animal protocols were approved by the Institutional Animal Care and Use Committee of Asahikawa Medical University. To evaluate in vivo degradability in two distinct biological environments, NF scaffolds were implanted either as sheet-type constructs in mice (subcutaneous model) or as small-caliber tubular grafts in rats (vascular model).

For the subcutaneous model, NF sheets were implanted into the dorsal subcutaneous tissue of male Jcl:ICR mice aged 6-8 weeks (Charles River). Under inhalation anesthesia with isoflurane, a midline dorsal incision was made, and four NF sheets were placed in the subcutaneous space and fixed with 7-0 polypropylene sutures (Prolene; Ethicon). The implants were harvested at 4, 8, 16, 24, and 48 weeks post-implantation. Retrieved samples were washed with phosphate-buffered saline (PBS) and used for SEC analysis. Samples used for SEM were additionally fixed overnight in 4% paraformaldehyde (PFA).

For the vascular model, NF grafts were implanted into the infrarenal abdominal aorta of male Wistar rats aged 8-10 weeks (Charles River) under isoflurane anesthesia. Following a midline laparotomy, the abdominal aorta was exposed, and the grafts were anastomosed using 9-0 polypropylene sutures (Prolene; Ethicon). Restoration of distal pulsation was confirmed before abdominal closure. The grafts were harvested at 4, 8, 16, 24, and 48 weeks after implantation. Samples used for SEC were collected after PBS perfusion via intracardiac infusion through the left ventricular apex under deep anesthesia and subsequent euthanasia. Samples used for SEM were perfusion-fixed with 4% PFA and then additionally immersion-fixed overnight in 4% PFA.

### 2.3. Size-Exclusion Chromatography (SEC)

SEC analysis was performed at Tosoh Analysis and Research Center Co., Ltd. using an HLC-8320GPC system (Tosoh Corporation) equipped with a TSKgel guard column HHR-H (6.0 mm I.D. × 4 cm) and TSKgel GMHHR-H columns (7.8 mm I.D. × 30 cm × 2). Chloroform (HPLC grade, containing amylene stabilizer; FUJIFILM Wako Pure Chemical Corporation) was used as the eluent at a flow rate of 1.0 mL/min, and the column temperature was maintained at 40 °C. Elution was monitored using a differential refractive index detector. For SEC analysis, only the soluble fraction of each retrieved sample was used. Samples were dissolved in the appropriate organic solvent at a concentration of 1 mg/mL overnight at room temperature, filtered through a 0.45 µm PTFE filter, and injected at a volume of 100 µL. Molecular weights were calculated as polystyrene-equivalent values using a calibration curve generated with standard polystyrene (Tosoh Corporation). Differential molecular weight distribution curves were used to evaluate changes in molecular weight profiles during degradation. Number-average molecular weight (Mn), weight-average molecular weight (Mw), and dispersity (Mw/Mn) were used for quantitative analysis.

### 2.4. Scanning Electron Microscopy (SEM)

Nanofiber morphology and surface topography were examined by SEM at the Toyama Industrial Technology Research and Development Center.

Paraformaldehyde-fixed specimens were dehydrated through a graded ethanol series (70%, 80%, 90%, and 95%; 30 min each). After dehydration, ethanol was replaced with t-butyl alcohol (1 mL, three times). The specimens were then freeze-dried using a freezer and a vacuum dryer (VD-400F; TAITEC Co.). After drying, the specimens were cut in liquid nitrogen using a trimming razor, mounted on aluminum stages, coated with gold using an ion sputter coater (JEC-550; JEOL Ltd.), and observed with a scanning electron microscope (JSM-6610LA; JEOL Ltd.).

### 2.5. Solid-State Nuclear Magnetic Resonance (NMR)

For solid-state NMR analysis, pre-implantation control scaffolds and the solvent-insoluble fragments remaining after SEC sample preparation of explanted scaffolds were examined. Prior to measurement, the samples were cut into pieces approximately 1 mm in width using scissors. Solid-state NMR measurements were performed on a Varian VNMRS-400 spectrometer. ^13^C cross-polarization magic-angle spinning (CP/MAS) NMR spectra were acquired at an observation frequency of 100.5 MHz with a contact time of 2 ms. A total of 8,192 scans were accumulated with a recycle delay of 3 s, and the MAS spinning rate was set at 7 kHz.

### 2.6. Statistical Analysis

Statistical comparisons were applied only to quantitative datasets obtained from replicate samples. Continuous variables are presented as mean ± standard deviation (SD). Differences among groups were assessed using one-way analysis of variance (ANOVA), assuming homogeneity of variance. When a significant overall difference was detected, multiple comparisons were performed using Tukey’s post hoc test. A p value of < 0.05 was considered statistically significant. All statistical analyses were performed using GraphPad Prism (Dotmatics).

## 3. Results

### 3.1. Environment-dependent degradation profiles of individual nanofiber scaffolds assessed by SEC

To characterize the environment-dependent in vivo degradation behavior of biodegradable NF scaffolds commonly used in vascular tissue-engineering studies—including PCL, PLA, PCL/PLA, and PGA—we implanted NF scaffolds into two distinct biological environments. Nanofiber sheets were implanted subcutaneously in mice, and small-caliber vascular grafts were implanted into the abdominal aorta of rats. The scaffolds were harvested at 4, 8, 16, 24, and 48 weeks after implantation, and degradation was evaluated by quantifying time-dependent changes in molecular weight using size-exclusion chromatography (SEC) as outlined in Figure 1.

**Figure 1.**
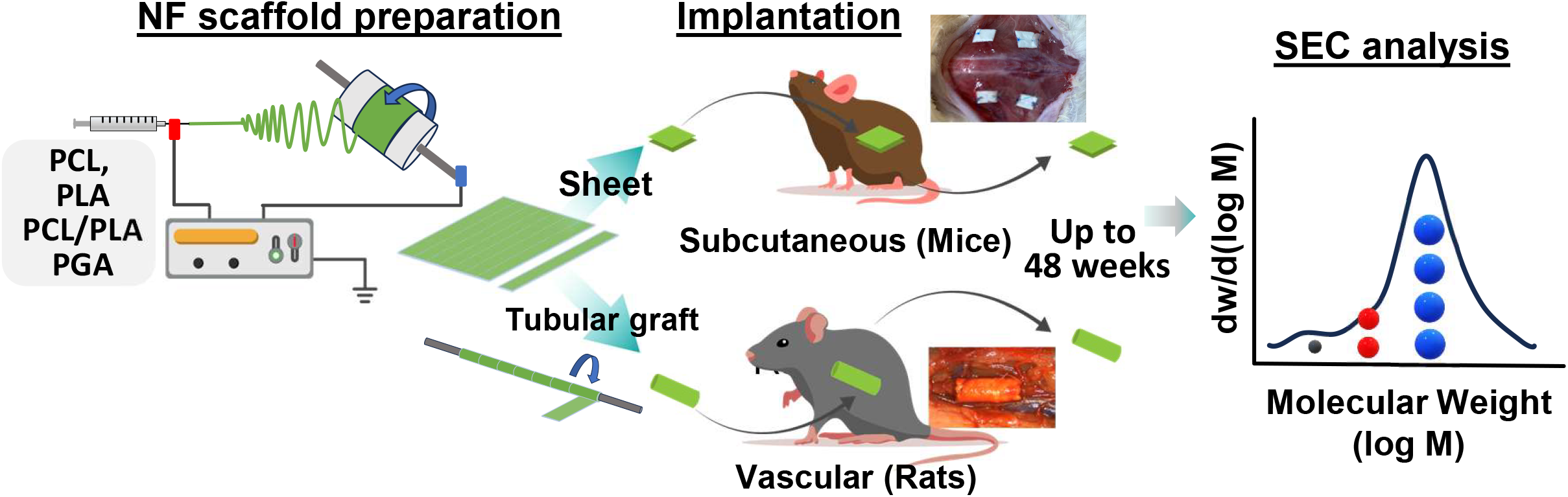
Overview of the experimental design for in vivo degradation analysis of electrospun nanofiber scaffolds. Electrospun nanofiber (NF) scaffolds composed of PCL, PLA, PCL/PLA, and PGA were prepared as sheet-type constructs and tubular grafts. Sheet-type scaffolds were implanted subcutaneously in mice, whereas tubular grafts were implanted into the abdominal aorta of rats. Scaffolds were harvested at 4, 8, 16, 24, and 48 weeks after implantation. In vivo degradation was quantitatively evaluated by SEC based on time-dependent changes in molecular weight distribution and SEC-derived molecular weight parameters.

#### PCL

The control PCL scaffold exhibited a single peak in the molecular weight distribution curve at approximately 5 MW (LogM) (Figure 2A). In both the subcutaneous and vascular environments, no marked shift of the peak position toward lower molecular weights was observed, whereas the peak height gradually decreased over time. In both environments, a new low–molecular-weight peak at approximately 3 MW (LogM) became detectable from 8 weeks post-implantation (Figure 2A). Notably, under the vascular environment, an additional intermediate–molecular-weight peak was detected between LogM 4.5 and 5.0, forming a gently broadened, skirt-like distribution (Figure 2A).

**Figure 2.**
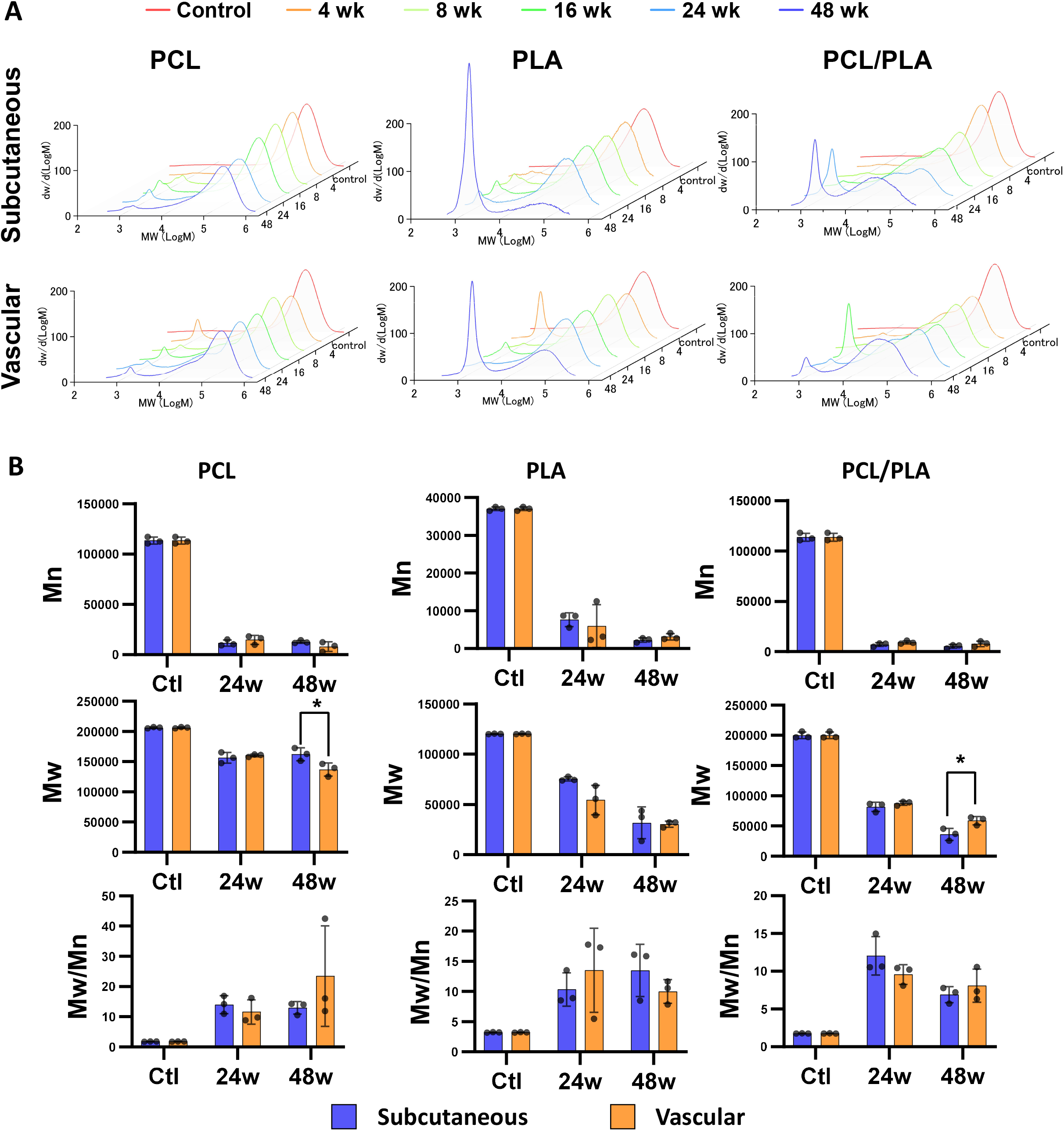
Size-exclusion chromatography (SEC) profiles and molecular weight parameters of electrospun nanofiber scaffolds in subcutaneous and vascular environments. **(A)** Time course of differential molecular weight distribution curves (dw/d(log M) vs. log M) of PCL, PLA, and PCL/PLA scaffolds after implantation. Upper and lower panels correspond to subcutaneous and vascular implantation, respectively. Curves are shown for control (red) and 4, 8, 16, 24, and 48 weeks post-implantation (orange, light green, green, light blue, and blue, respectively). **(B)** Quantitative SEC-derived molecular weight parameters (Mn, Mw, and Mw/Mn) for PCL, PLA, and PCL/PLA at control (Ctl), 24 weeks (24w), and 48 weeks (48w). Blue and orange bars indicate subcutaneous and vascular samples, respectively. Dots represent individual samples, and bars show mean ± SD. *p<0.05 between environments at the indicated time point.

Quantitative SEC analysis demonstrated significant decreases in both Mn and Mw in the subcutaneous and vascular environments by 48 weeks. Mn decreased rapidly, reflecting the generation of low–molecular-weight species during degradation, whereas Mw exhibited a more gradual decline, indicating progressive degradation of high–molecular-weight polymer chains. Accordingly, the Mw/Mn ratio increased progressively over time, indicating a broadening of the molecular weight distribution during degradation. Comparison between environments revealed no significant difference in Mn at either 24 or 48 weeks. In contrast, Mw was significantly lower in the vascular environment than in the subcutaneous environment at 48 weeks (vascular: 137,070 ± 10,937 vs. subcutaneous: 162,320 ± 10,742, p = 0.046; Figure 2B), whereas no difference was observed at 24 weeks. These data indicate accelerated degradation of the high–molecular-weight fraction under vascular conditions during the late phase.

#### PLA

In the subcutaneous environment, PLA sheets remained macroscopically identifiable at 36 weeks after implantation (Supplementary Figure S1A). By 48 weeks, some PLA sheets became visually unidentifiable, indicating near-complete degradation, and could not be retrieved; therefore, only the remaining sheets were subjected to SEC analysis. In contrast, in the vascular environment, PLA scaffolds remained macroscopically intact without obvious morphological changes up to 48 weeks (Supplementary Figure S1B).

SEC profiles demonstrated a slight leftward shift of the main molecular weight peak at approximately log M = 5.0 in both environments over time (Figure 2A). A low–molecular-weight peak at approximately log M = 3 became detectable from 4 weeks and increased markedly by 48 weeks. The reduction in the height of the high–molecular-weight peak was more pronounced in the subcutaneous environment at 48 weeks. Quantitative SEC analysis indicated early formation of low–molecular-weight species (Mn) by 24 weeks, followed by a progressive reduction of the high–molecular-weight fraction (Mw) and broadening of the molecular weight distribution (Mw/Mn) over time.

Despite the marked macroscopic difference at 48 weeks, SEC-based molecular weight parameters did not show clear differences between the subcutaneous and vascular environments. This may be partly attributable to the subcutaneous 48-week group, in which only the residual scaffold material was available for analysis; thus, the quantified values primarily reflect the properties of the residual fraction rather than the entire implanted scaffold.

Taken together, these results indicate that PLA nanofibers exhibited a markedly environment-dependent degradation behavior.

#### PCL/PLA

The control PCL/PLA samples showed a single main peak at approximately log M = 5.0. In both subcutaneous and vascular environments, an additional intermediate–molecular-weight peak emerged between log M 4.5 and 5.0 by 8 weeks post-implantation, resulting in a bimodal distribution pattern (Figure 2A). By 48 weeks, the original peak at approximately log M = 5.0 was markedly reduced, whereas the intermediate peak became dominant; however, the peak position of this predominant intermediate fraction differed between environments. In the subcutaneous environment, the dominant peak shifted toward the lower-molecular-weight side, whereas in the vascular environment it tended to remain at a relatively higher molecular weight (Figure 2A).

This trend was further supported by quantitative SEC analysis: Mw at 48 weeks was significantly lower in subcutaneously implanted samples than in vascularly implanted samples (subcutaneous: 36,036 ± 10,053 vs. vascular: 59,002 ± 6,983, p = 0.031; Figure 2B). Taken together, these results indicate that the PCL/PLA blend underwent a time-dependent shift in its molecular weight distribution, with more pronounced degradation of the high–molecular-weight fraction in the subcutaneous environment during the late phase.

#### PGA

PGA nanofibers showed rapid degradation in vivo. In the vascular implantation model, PGA grafts lost structural integrity and ruptured within 1 week, limiting further evaluation in the vascular environment. Therefore, PGA degradation was assessed only in the subcutaneous model, where samples could be retrieved up to 2 weeks post-implantation (Supplementary Figure S2). SEC profiles of PGA in the subcutaneous environment showed marked time-dependent changes in the molecular weight distribution. By 1 week, the original high–molecular-weight peak was substantially diminished. At 2 weeks, an additional peak appeared at a slightly higher molecular weight than the original peak (Supplementary Figure S2A). Quantitative SEC analysis showed a transient decrease in both Mn and Mw at 1 week, followed by recovery to levels comparable to the control by 2 weeks (Supplementary Figure S2B). Meanwhile, the Mw/Mn ratio increased over time, indicating progressive broadening of the molecular weight distribution during degradation (Supplementary Figure S2B). Together with the detection of an additional higher-molecular-weight peak at 2 weeks (Supplementary Figure S2A), these findings suggest a transient, non-monotonic change in the molecular weight distribution during the early phase of PGA degradation in vivo.

Because PGA could not be evaluated beyond 2 weeks and vascular data were unavailable due to early graft rupture, PGA was not included in subsequent comparative analyses (Sections 3.2–3.4).

### 3.2. Comparative analysis of relative molecular weight changes among nanofiber materials

Next, to compare differences in degradation behavior among materials, relative changes in SEC-derived molecular weight parameters were analyzed. All relative values were expressed as percentages relative to the corresponding control values (Figure 3). In the subcutaneous environment, Mn decreased markedly by 24 and 48 weeks across all materials, consistent with the generation of low–molecular-weight chains during degradation (Figure 3A). In contrast, the extent of Mw reduction differed substantially among materials. At 24 weeks, the relative Mw values were approximately 80%, 60%, and 40% for PCL, PLA, and PCL/PLA, respectively, and the differences among materials were statistically significant. At 48 weeks, the relative Mw values were approximately 80%, 20%, and 10%, respectively, with Mw values for PLA and PCL/PLA being significantly lower than that for PCL. These findings indicate that degradation of the high–molecular-weight fraction proceeds in a material-dependent manner. PCL retained a relatively high Mw up to 48 weeks, suggesting prolonged persistence of high–molecular-weight chains. In contrast, PLA exhibited a pronounced decrease in Mw from 24 to 48 weeks, indicating that degradation of the high–molecular-weight fraction became more prominent during the later phase. The PCL/PLA blend showed the largest reduction in Mw already at 24 weeks, suggesting earlier degradation of high–molecular-weight components compared with PCL and PLA. To assess degradation-associated heterogeneity, Mw/Mn (normalized to the control) was evaluated as an index of the breadth of the molecular weight distribution. At both 24 and 48 weeks, Mw/Mn was significantly higher in PCL than in PLA and PCL/PLA. This indicates greater molecular weight heterogeneity in PCL, consistent with the coexistence of low–molecular-weight species (lower Mn) and a relatively preserved high–molecular-weight fraction (higher Mw) during degradation.

**Figure 3.**
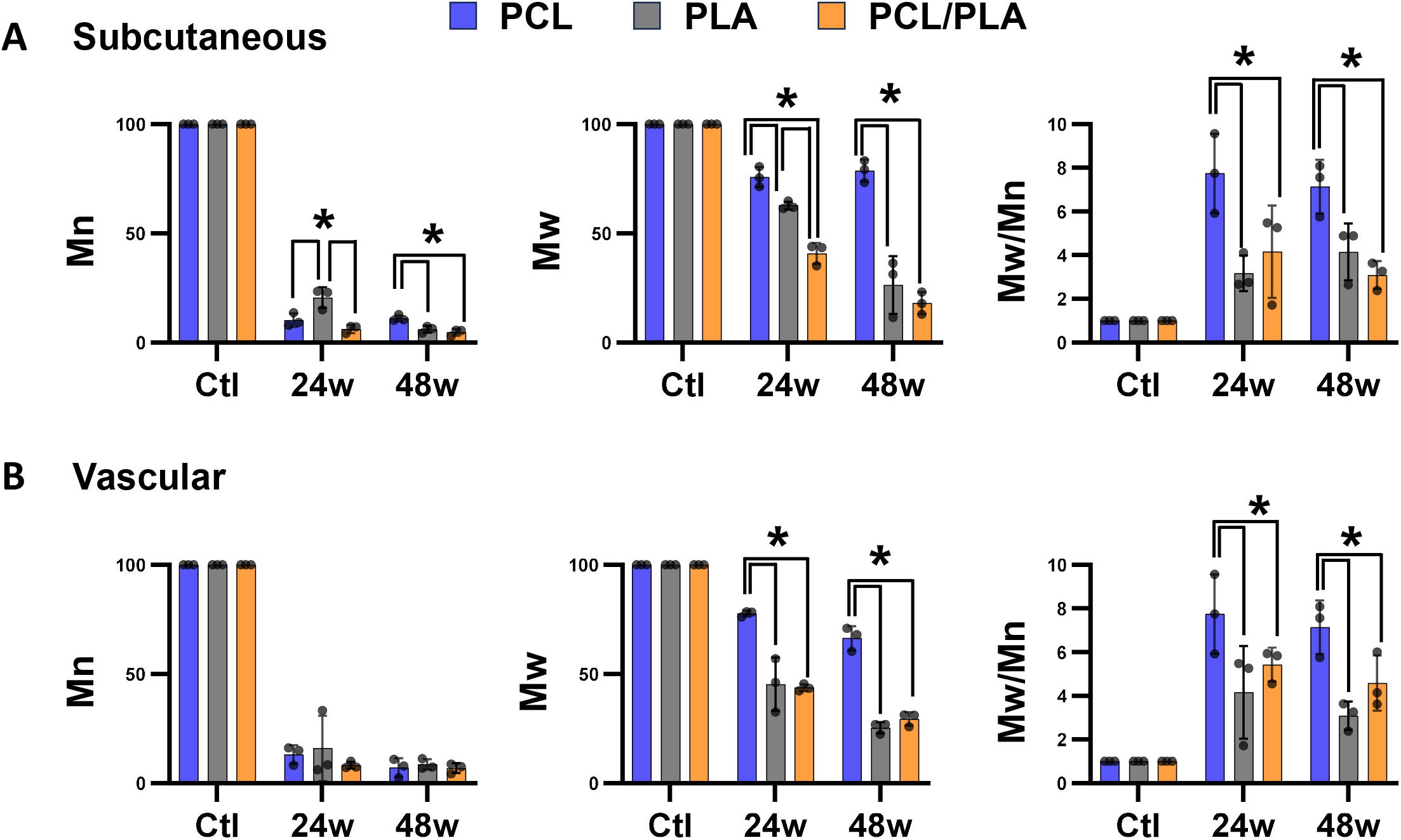
Relative changes in SEC-derived molecular weight parameters among nanofiber materials in subcutaneous and vascular environments. Relative values of number-average molecular weight (Mn) and weight-average molecular weight (Mw) are presented as percentages of the corresponding control (Ctl) values, and dispersity (Mw/Mn) is shown normalized to the control (= 1). **(A)** Subcutaneous implantation model. Relative Mn, relative Mw, and normalized Mw/Mn for PCL, PLA, and PCL/PLA at 24 and 48 weeks post-implantation. **(B)** Vascular implantation model. Relative Mn, relative Mw, and normalized Mw/Mn for PCL, PLA, and PCL/PLA at 24 and 48 weeks post-implantation. Data are shown as mean ± SD; dots represent individual samples. *p < 0.05.

In the vascular environment, Mn also decreased markedly by 24 and 48 weeks across all materials (Figure 3B). At 24 weeks, the relative Mw values were approximately 80%, 45%, and 45% for PCL, PLA, and PCL/PLA, respectively, and at 48 weeks were approximately 70%, 25%, and 30%. At both time points, Mw values for PLA and PCL/PLA were significantly lower than that for PCL. Similar to the subcutaneous environment, Mw/Mn values were consistently higher for PCL than for the other materials in the vascular environment (Figure 3A and 3B), indicating preferential retention of high–molecular-weight chains during degradation. In contrast, PLA exhibited an increase in Mw/Mn at 24 weeks relative to the control, followed by a decrease toward 48 weeks. This pattern suggests an initial broadening of the molecular weight distribution during degradation, followed by narrowing as high–molecular-weight fractions were progressively lost. These findings indicate that degradation pathways and molecular weight distribution dynamics differ among nanofiber materials.

Overall, the trends observed in the vascular environment were largely consistent with those in the subcutaneous environment: PCL retained a larger fraction of high–molecular-weight chains, whereas PLA and PCL/PLA exhibited more pronounced degradation of high–molecular-weight fractions by 48 weeks. Notably, PLA showed a more pronounced reduction in Mw at 24 weeks in the vascular environment compared with the subcutaneous environment.

### 3.3. Ultrastructural changes in nanofiber scaffolds during in vivo degradation evaluated by electron microscopy

To evaluate changes in nanofiber ultrastructure during the in vivo degradation process, explanted scaffolds were examined by SEM. In the pre-implantation controls, the scaffolds showed a homogeneous fibrous cross-sectional morphology with clearly identifiable nanofiber profiles across all materials (PCL, PLA, and PCL/PLA) and scaffold formats (subcutaneous sheets and vascular grafts). After implantation, this fibrous appearance began to become less apparent as early as 8 weeks, with cross sections increasingly showing smoother chunk-like areas rather than discrete fiber profiles (Figure 4). This progressive loss of fibrous architecture was observed across all materials and became most pronounced at 48 weeks; the change was most obvious in PCL/PLA. (Figure 4). Collectively, these observations indicate a time-dependent loss of the nanofiber architecture in vivo, even when the scaffold remained macroscopically identifiable up to 48 weeks.

**Figure 4.**
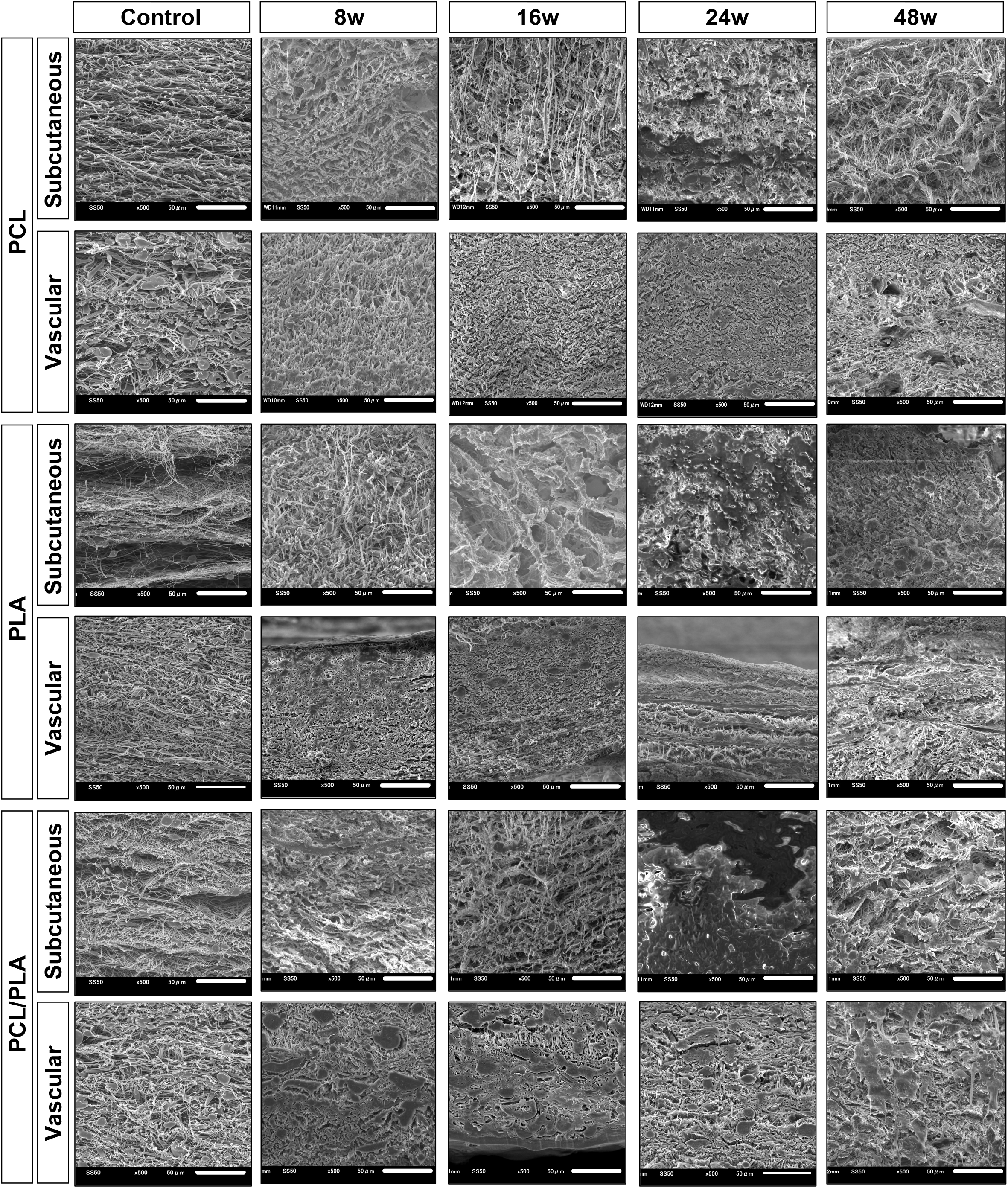
Scanning electron microscopy (SEM) images of nanofiber scaffold microstructure. Representative SEM images of explanted PCL, PLA, and PCL/PLA scaffolds harvested at 8, 16, 24, and 48 weeks post-implantation are shown for the subcutaneous sheet model and the vascular graft model, as indicated. Control is the pre-implantation scaffold. All images were acquired at 500× magnification. Scale bar = 50 µm.

### 3.4. Characterization of the insoluble fraction of explanted nanofiber scaffolds by solid-state NMR

During SEC sample preparation, nanofiber scaffolds were dissolved in organic solvents and only the soluble fraction was subjected to analysis. While pre-implantation control nanofibers dissolved completely, explanted scaffolds consistently left an insoluble residue after solvent treatment (Figure 5A). To characterize the chemical composition of this insoluble fraction, solid-state ^13^C NMR spectra were acquired using the CPMAS method. Analysis was performed on control scaffolds and subcutaneous PCL scaffolds retrieved at 24 weeks as a representative sample. In both control and explanted samples, characteristic PCL resonances were observed at 25–34 ppm (CH_2_), ∼65 ppm (O–CH_2_), and ∼174 ppm (C=O), confirming the presence of PCL-derived components in the insoluble material (Figure 5B). In addition to the PCL signals, explanted PCL scaffold sample exhibited several additional resonances, including peaks in the 40–80 ppm region and at 166–190 ppm, consistent with amide-associated carbons and amide C=O, respectively (Figure 5B). Signals in the 124–134 ppm region suggested the presence of aromatic carbon–containing components, although their identity could not be determined. Similarly, explanted PLA and PCL/PLA samples also showed additional signals consistent with amide-associated carbons and amide C=O. Collectively, these results indicate that the solvent-insoluble fraction observed after implantation contains polymer-derived components together with additional signals consistent with biological constituents.

**Figure 5.**
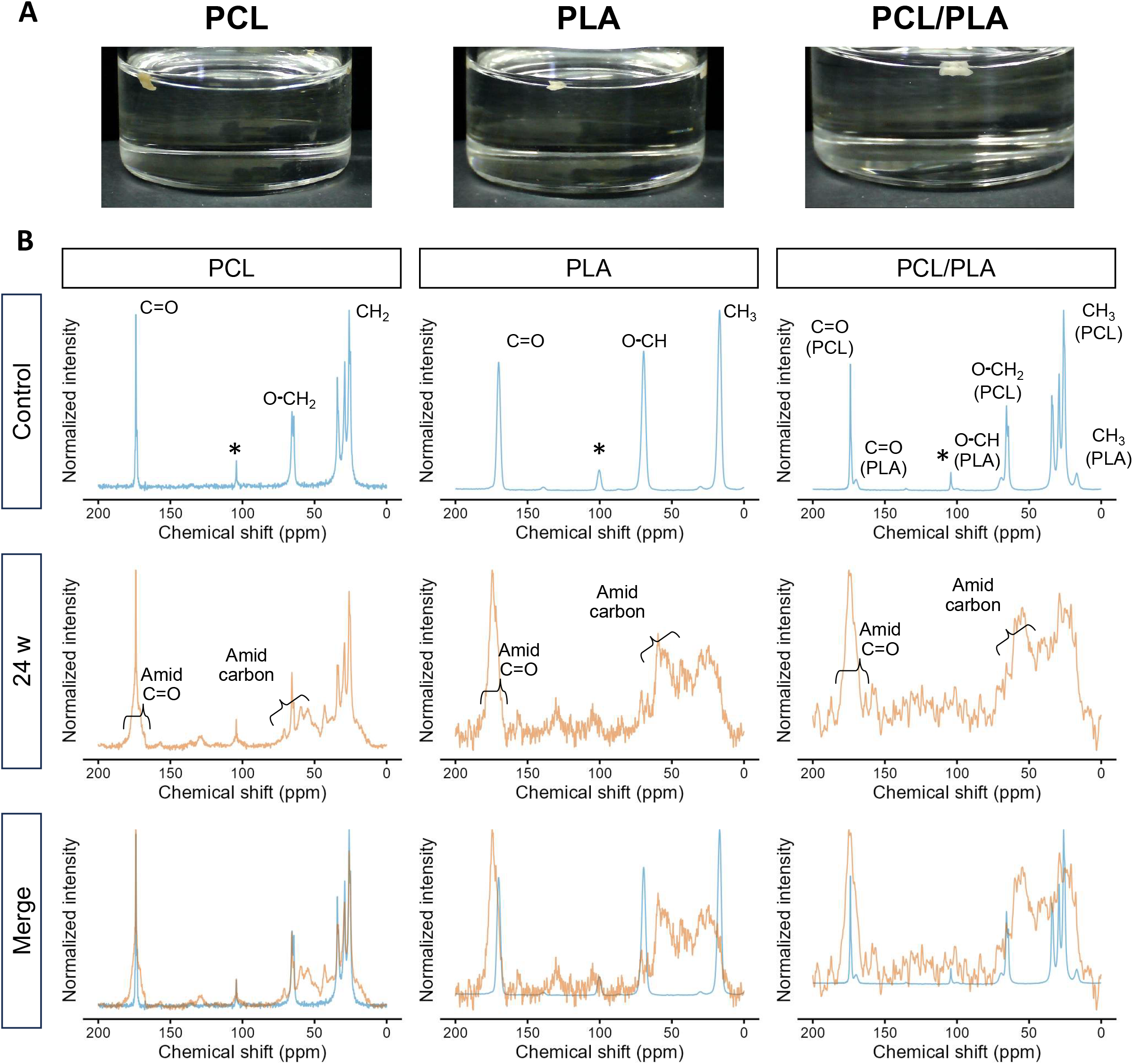
Solid-state ^13^C NMR analysis of explanted nanofiber scaffolds. **(A)** Representative images of the insoluble residues remaining after organic solvent treatment during SEC sample preparation. PCL, PLA, and PCL/PLA scaffolds were explanted at 24 weeks after implantation. **(B)** Solid-state ^13^C CPMAS NMR spectra of the corresponding scaffolds. For each material, spectra are shown for the pre-implantation control (top), the subcutaneous sample retrieved at 24 weeks (middle), and the overlay of the two spectra (bottom). Characteristic polymer-derived resonances are annotated for each material in the control spectrum (top). Additional signals observed in the explanted samples are indicated as amide carbon and amide C=O in the 24-week spectrum (middle). The asterisk indicates a spinning side band.

## 4. Discussion

In this study, we quantitatively characterized the in vivo degradation of electrospun biodegradable nanofiber scaffolds (PCL, PLA, PCL/PLA, and PGA). Time-dependent changes in molecular weight were evaluated using SEC. This analysis showed that degradation behavior differed not only among materials but also between subcutaneous and vascular environments. In addition, scaffold degradation was accompanied by progressive loss of nanofiber ultrastructure, as well as altered solubility and chemical composition. These findings suggest that in vivo nanofiber degradation is a complex process involving microstructural alteration and interaction with the biological environment.

### 4.1 Implications of Nanofiber Degradation for Vascular Graft Design

The present study provides a comparative assessment of the in vivo degradation behavior of biodegradable electrospun nanofiber scaffolds that have been widely investigated for small-caliber vascular grafts. Among the materials evaluated, PGA has previously been reported to exhibit rapid degradation and limited durability when used as a standalone scaffold [20], and this observation was reproduced in the present study, as evidenced by early loss of structural integrity and graft rupture within 1 week in the vascular environment. This behavior was also recapitulated by the rapid reduction in molecular weight observed in the subcutaneous model (Figure S2), supporting the utility of molecular weight change as a quantitative indicator of in vivo.

In contrast, PCL, PLA, and PCL/PLA scaffolds remained macroscopically intact for up to 48 weeks in the vascular environment, while showing distinct degradation profiles. PCL maintained the highest Mw in both subcutaneous and vascular environments, whereas PLA and PCL/PLA showed greater reductions in Mw over time (Figure 3). These findings are consistent with the general understanding that PCL degrades slowly, whereas incorporation of PLA accelerates scaffold degradation [21]. The slow degradation of PCL suggests its suitability when prolonged mechanical support is required, whereas the faster degradation of PLA and PCL/PLA may favor earlier scaffold degradation and remodeling. Comparative evaluation of degradation profiles is therefore essential for rational material selection in biodegradable vascular grafts. Further studies are needed to clarify how molecular weight changes relate to vascular compliance and tissue regeneration in vivo[16].

### 4.2 Environmental Dependence of Nanofiber Degradation

Degradation of nanofiber scaffolds was evaluated in both subcutaneous and vascular environments in this study. Although broadly similar time-dependent patterns were observed for all tested materials in the two settings, the extent of degradation differed between environments, particularly in the later phase. In PCL, the high–molecular-weight fraction was significantly better preserved in the subcutaneous environment than in the vascular environment, whereas PCL/PLA showed the opposite pattern at 48 weeks after implantation (Figure 2B, Mw). This difference was most evident in PLA, which underwent near-complete degradation in the subcutaneous environment by 48 weeks, but not in the vascular environment at the same time point.

PLA degradation has been reported to be influenced by water availability, local pH, and autocatalytic effects associated with the accumulation of degradation products [22–24], which may help explain the environmental difference observed in the present study. In the subcutaneous environment, degradation products are more likely to remain around the implant, which may promote local acidification and accelerate subsequent degradation. In contrast, in the vascular environment, degradation products may be continuously removed by blood flow, thereby reducing local accumulation and limiting autocatalytic effects [24]. Although further studies are needed to clarify the detailed mechanisms underlying these differences, the present findings have important implications for experimental design. Subcutaneous implantation is a convenient in vivo model; however, the present results indicate that degradability assessed under subcutaneous conditions cannot be directly extrapolated to vascular applications. For vascular graft development, evaluation under vascular conditions is therefore essential.

### 4.3 Complexity of In Vivo Nanofiber Degradation

As shown in Figure 5, explanted scaffolds consistently left an insoluble fraction after organic solvent treatment during SEC sample preparation, whereas pre-implantation control scaffolds dissolved completely under the same conditions. Because biodegradable vascular graft scaffolds are generally expected to be progressively replaced by host tissue after implantation, one possible interpretation was that this insoluble fraction primarily represented regenerated host tissue. However, solid-state ^13^C NMR demonstrated that the insoluble fraction contained not only signals derived from biological constituents, but also polymer-derived resonances. These findings indicate that the insoluble fraction cannot be explained solely as host-derived tissue that remained insoluble after solvent treatment. Rather, they suggest that at least part of the implanted nanofiber-derived material had become resistant to solubilization under the SEC preparation conditions.

Although the mechanism underlying this change remains unclear, several possibilities may be considered. One is association of the degrading scaffold with biological constituents, leading to the formation of insoluble polymer–biological complexes [25,26]. Another is physicochemical change in the scaffold itself, such as increased crystallinity or molecular reorganization during degradation [27]. Further studies will be required to distinguish between these possibilities. Nevertheless, the present findings indicate that the implanted scaffold may no longer remain chemically and physically equivalent to the original scaffold, even when scaffold-derived components are still present [28].

The SEM findings are consistent with this interpretation. Even when scaffolds remained macroscopically identifiable, their nanofibrous architecture progressively changed over time, with loss of distinct fiber profiles. Together with the NMR results, these observations suggest that degradation of implanted nanofiber scaffolds proceeds as a multilevel process involving not only molecular weight reduction, but also altered solubility, chemical compositional change, and loss of fibrous ultrastructure. Further studies will be required to determine the precise mechanisms responsible for these changes, but the present results indicate that in vivo nanofiber degradation is not a simple process of hydrolysis alone and is likely shaped by complex interactions with the biological environment.

## 5. Conclusion

In this study, we characterized in vivo molecular weight changes in biodegradable electrospun nanofiber scaffolds and demonstrated that degradation behavior differs among materials and is influenced by the implantation environment. In addition to molecular weight reduction, scaffold degradation was associated with loss of fibrous ultrastructure and changes in chemical composition, indicating that in vivo degradation is more complex than simple hydrolysis alone. These findings provide a framework for evaluating scaffold degradability and offer a new perspective on the design of biodegradable nanofiber vascular grafts.

## Abbreviations

NF: nanofiber
PCL: polycaprolactone
PLA: poly(D,L-lactide)
PGA: polyglycolic acid
PCL/PLA: polycaprolactone/poly(D,L-lactide) blend
SEC: size-exclusion chromatography
Mn: number-average molecular weight
Mw: weight-average molecular weight
SEM: scanning electron microscopy
NMR: nuclear magnetic resonance

## Acknowledgments

We thank the staff of the Tosoh Analysis Center for their technical support and assistance with the SEC and solid-state NMR measurements.

## Data statement

The datasets generated and analyzed during the current study are available from the corresponding author upon reasonable request.

## Author contributions (CRediT)

Shingo Kunioka: Conceptualization, Data curation, Formal analysis, Methodology, Validation, Visualization, Writing – original draft.

Takumi Yoshida: Investigation, Resources.

Daisuke Naruse: Investigation, Resources.

Yuki Setogawa: Investigation.

Hiroyuki Miyamoto: Investigation.

Ryohei Ushioda: Investigation.

Yuta Kikuchi: Investigation, Supervision.

Masahiro Tsutsui: Investigation, Supervision.

Hiroyuki Kamiya: Conceptualization, Supervision, Writing – review and editing.

Kyohei Oyama: Conceptualization, Supervision, Writing – review and editing.

---

**Supplemental Figure S1.**
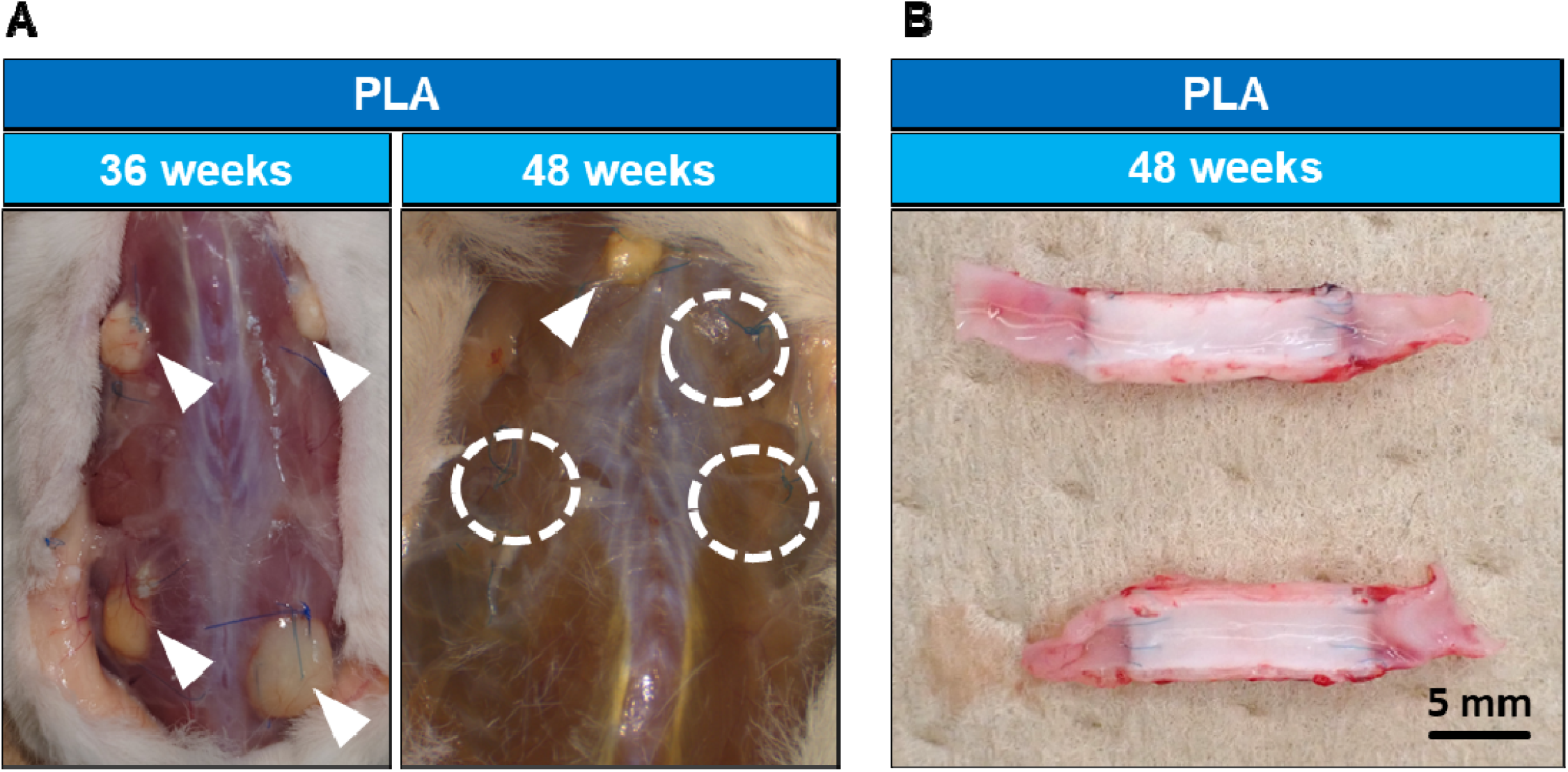
Macroscopic appearance of implanted scaffolds at explantation. **(A)** Representative images of PLA sheets at 36 and 48 weeks after subcutaneous implantation. White arrowheads indicate the implanted PLA sheets. Dashed circles indicate the implantation sites where the PLA sheets had been placed; these sites were identified using the fixation sutures as landmarks. At 36 weeks, all implanted PLA sheets were visually identifiable, whereas at 48 weeks, some PLA sheets were no longer visually detectable. **(B)** Representative image of a PLA vascular graft at 48 weeks after vascular implantation. All grafts remained visually identifiable up to 48 weeks.

**Supplemental Figure S2.**
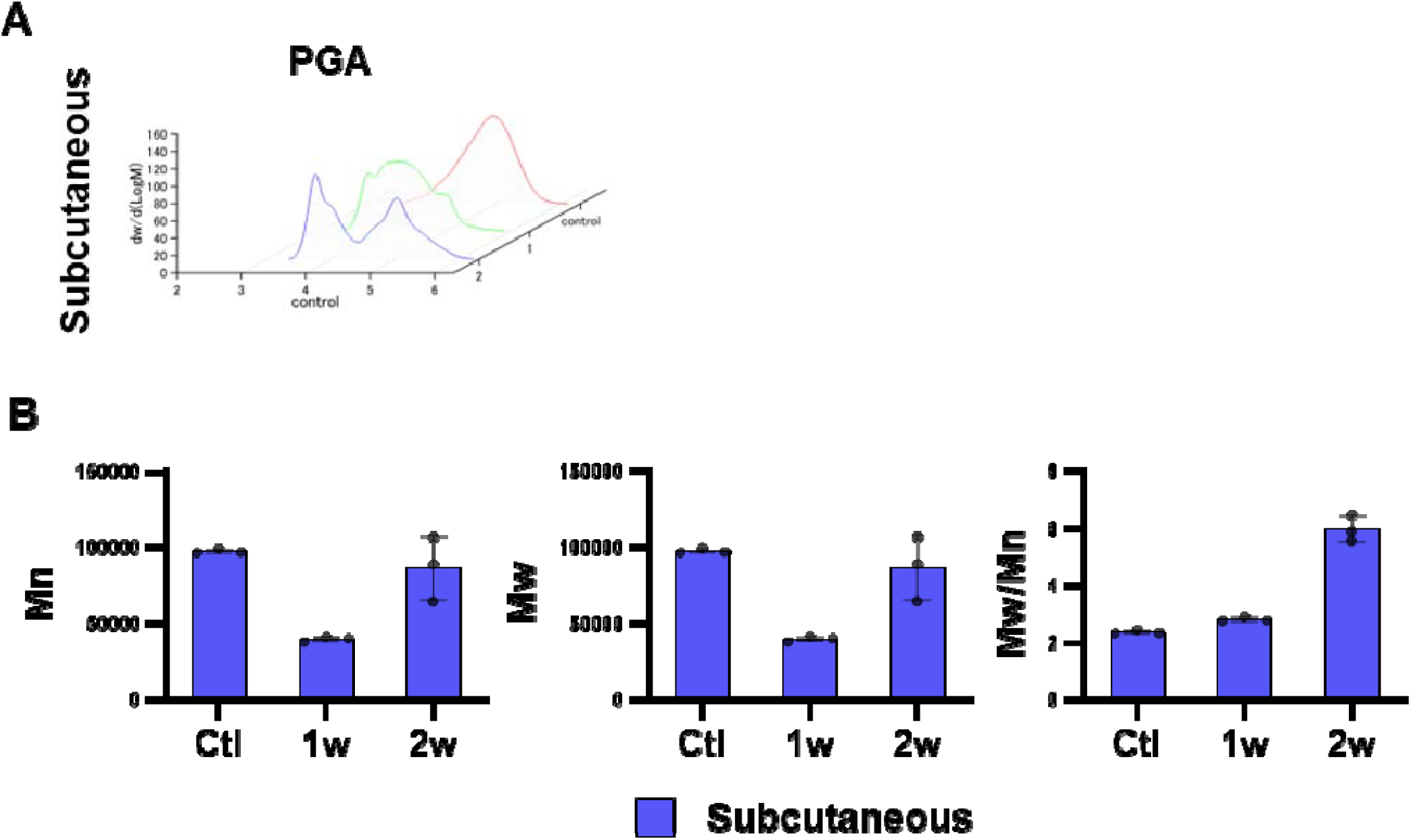
SEC analysis of PGA nanofiber scaffold in the subcutaneous environment. (A) Differential molecular weight distribution curves of explanted PGA scaffolds retrieved from the subcutaneous implantation model (control, 1 week, and 2 weeks). The original high–molecular-weight peak was markedly reduced at 1 week, and an additional higher-molecular-weight peak was detected at 2 weeks. (B) Quantitative SEC-derived parameters (Mn, Mw, and Mw/Mn) at each time point. Mn and Mw showed a transient decrease at 1 week followed by recovery to levels comparable to the control at 2 weeks, whereas Mw/Mn increased over time. Values are presented as mean ± SD.

